# Tactile Pressure Evokes a Biphasic BOLD Response in Ipsilateral Primary Somatosensory Cortex

**DOI:** 10.1101/2025.08.11.669660

**Authors:** Anna C. Feldbush, Nahid Kalantaryardebily, Neha A. Reddy, Rebecca Faubion-Trejo, Jeff Soldate, Jonathan Lisinski, Molly G. Bright, Netta Gurari, Stephen M. LaConte

## Abstract

Tactile perception is fundamental to how we engage with and interpret our surroundings. While the contralateral primary somatosensory cortex (S1) is thought to be generally responsible for processing tactile information, simultaneous responses in the ipsi-lateral hemisphere have been observed. This work aims to characterize the blood-oxygen-level-dependent (BOLD) response pattern of ipsilateral S1 to unilateral tactile stimuli. In this study, tactile stimuli were applied using a custom pneumatically actuated stimulator designed and built in house (KalantaryArdebily et al., 2024). Three stimulus force levels were applied in an event-related design. As expected we observed an apparent difference in BOLD responses between the contralateral and ipsilateral hemispheres. We found differing response patterns between the Brodmann’s areas (BA) within ipsilateral S1. Ipsilateral BA2 had a positive BOLD response similar to that of the contralateral hemisphere. In contrast, ipsilateral BA1 and BA3b appeared to have a biphasic response. That is, those regions had an initial negative response, followed by a secondary positive one. Using multi-echo analysis, we verified that this biphasic response pattern is BOLD-related. In addition, the secondary (positive) phase was more sensitive to tactile stimulus force than the initial “negative” BOLD phase. The ipsilateral BA1 and BA3b responses support a previous hypothesis of bilateral sensory gating (Blatow et al., 2007; Chung et al., 2014; Hämäläinen et al., 2000; Kastrup et al., 2008; Klingner et al., 2016; Schäfer et al., 2012; Tamè et al., 2015). This study extends previous research reporting negative responses in ipsilateral S1, suggesting that the response is BOLD-related, biphasic, and that the previously overlooked secondary positive phase is actually more sensitive to stimulus intensity.

## 1 Introduction

While it is well known that the contralateral primary somatosensory cortex (S1) responds to unilateral tactile stimuli, our understanding of ipsilateral responses has evolved significantly over the last three decades. Clarifying the functional role of ipsilateral S1 may yield valuable insights into a phenomenon that is both poorly understood and often overlooked. Beyond its implications for interhemispheric integration and distributed neural processing, ipsilateral responses may provide crucial insights into compensatory function after peripheral nerve damage (Fornander et al., 2016) and central neurological injury from stroke and traumatic brain injury. Early studies in neurosurgical patients first suggested that median nerve stimuli could evoke ipsilateral electrical responses (Allison et al., 1989; Noachtar et al., 1997). The occurrence and characterization of ipsilateral S1 responses has since been studied with EEG (Sutherland & Tang, 2006), MEG (Kanno et al., 2003; Korvenoja et al., 1995; Korvenoja et al., 1999; Lipton et al., 2006; Schnitzler et al., 1995), and fMRI (Blatow et al., 2007; Eickhoff et al., 2008; Korvenoja et al., 1999; Nihashi et al., 2005; Stringer et al., 2014). However, ipsilateral findings have been less robust and less reproducible than contralateral S1 responses. Moreover, the interpretation of ipsilateral responses in fMRI studies has been complicated by the observation that median nerve stimuli tend to evoke negative blood-oxygen-level-dependent (BOLD) responses in ipsilateral S1 (Blatow et al., 2007; Eickhoff et al., 2008; Hlushchuk & Hari, 2006; Kanno et al., 2003; Klingner, Huonker, et al., 2011; Klingner et al., 2014; Lamp et al., 2019; Lipton et al., 2006; Nihashi et al., 2005; Schäfer et al., 2012; Stringer et al., 2014). Similar to negative BOLD phenomena reported in the visual cortex (Amedi et al., 2005; Shmuel et al., 2006), definitive mechanistic interpretations of negative BOLD remains elusive (Gouws et al., 2014; Klingner et al., 2014; Nelson & Mayhew, 2025; Schäfer et al., 2012).

To address these challenges in interpreting negative responses in ipsilateral S1, the current study focuses on two important facets of BOLD measurements. First, we employed a multi-echo echo-planar imaging (EPI) sequence to assess the echo-time (TE) and T2* dependence of the ipsilateral response. This enabled us to directly evaluate whether observed ipsilateral signals are indeed related to the BOLD mechanism. Second, we examined the hemodynamic response function (HRF) associated with negative BOLD signals, leveraging the fact that our stimulus duration was brief compared to most previous studies. Early multi-echo work in by Peltier and Noll, 2002, corroborated resting state data as neurovascularly coupled (BOLD-related) as opposed to arising from non-neuronal artifacts such as vascular symmetry, motion, physiological noise, and scanner instability. Further work has built on echo-time dependence as a hallmark characteristic of the BOLD response (Devi et al., 2022; DuPre et al., 2021; Havlicek et al., 2017; Kundu et al., 2012; N. Li & Jasanoff, 2020). Generally, this approach leverages the TE-dependence and decay properties of the signal to validate whether a given fluctuation arises from changes in T2*, which is the fundamental basis of BOLD contrast. While this approach has supported claims of positive BOLD-related signals, it has rarely if ever been applied to the study of negative BOLD responses.

Beyond T2* measurements, improved characterization of the HRF is critical for interpreting the neural and physiological underpinnings of BOLD signals. Within rather broad ranges of stimulus parameters, the BOLD signal can be approximated as the stimulus timing convolved with an HRF (Glover, 1999; Henson & Friston, 2007), although it is known that distinct neural or vascular mechanisms are not always well captured by canonical HRF models (Chen et al., 2023). Nevertheless, HRF convolution modeling remains reasonably accurate, even in the cases where these known limitations play a role (Boynton et al., 1996; Vazquez & Noll, 1998). Importantly, most studies reporting negative BOLD responses in ipsilateral S1 have used block designs, likely to enhance detection sensitivity given that ipsilateral responses are typically weaker and less reliable than contralateral ones. However, upon reviewing this literature, we observed that many of the reported negative responses deviate from the expected shape predicted by standard HRF convolution models (eg. Hlushchuk and Hari, 2006; Klingner, Huonker, et al., 2011). Specifically, given a block design stimulus of duration S and an HRF of duration H, the expected BOLD response (B) should equal S + H - 1 for sampled, discrete signals. Thus, a negative BOLD HRF under a block-design stimulus should produce a negative signal convolved beyond the duration of the block. Instead, previous reports frequently show negative BOLD responses lasting for only a portion of the stimulus duration (de la Rosa et al., 2021; Hlushchuk & Hari, 2006; Kastrup et al., 2008; Shmuel et al., 2006). We hypothesized two explanations for this discrepancy. One is that the negative BOLD response fundamentally violates linearity assumptions, suggesting mechanisms independent of neurovascular coupling. A second explanation would be that the hemodynamic response is biphasic with an initial negative phase followed by a positive rebound. Such a biphasic HRF could account for the short negative signal phase reported in previous studies where the negative response ends before the stimulus block itself concludes. Based on theoretical considerations and preliminary observations, we favored the biphasic hypothesis, suspecting it had been overlooked in earlier studies, potentially due to their long stimulus durations. To test this explicitly, we used a pneumatic stimulation device capable of delivering robust yet brief (*<*1 s duration) tactile stimuli to the tip of the right index finger at three distinct pressure levels (KalantaryArdebily et al., 2024). Given that our stimulus duration was relatively brief (S *<* H), we predicted that the negative BOLD response would appear after stimulation. Finally, informed by foundational studies on negative BOLD in S1 (Blatow et al., 2007; Eickhoff et al., 2008; Hlushchuk & Hari, 2006; Kanno et al., 2003; Klingner, Huonker, et al., 2011; Klingner et al., 2014; Lamp et al., 2019; Lipton et al., 2006; Nihashi et al., 2005; Schäfer et al., 2012; Stringer et al., 2014), we performed region of interest (ROI) analyses specifically targeting ipsilateral S1.

## 2 Methods

### 2.1 Participants

We collected fMRI data from 14 right-hand dominant individuals (male:7 female:7), as classified by the Edinburgh Handedness Inventory (Robinson, 2013). Participants were aged 18-23 (21±1.8 years). Additionally, participants reported no history of neurological or musculoskeletal injuries that could affect tactile signaling. Data collection was approved by Virginia Polytechnic University’s Institutional Review Board. All participants provided informed consent before experimentation and were aware of the study design and the nature of the tactile stimuli we provided. Participants were instructed that they could withdraw consent at any time.

### 2.2 Tactile Stimulation

We applied tactile stimuli to the tip of the right index finger using a custom-designed pneumatic actuator (Fig. 1, KalantaryArdebily et al., 2024). This device provides sustained tactile stimuli through the inflation of a small-radius balloon (7.5 mm). Our experiment used three different pressure levels (Low:88.7 kPa, Medium:101.3 kPa, High:116 kPa), which were sustained for approximately 0.67 s. Stimulus magnitude was controlled by changing the pressure inside the balloon, and thus the pressure applied to the finger. Across the low, medium, and high pressure levels there was a negligible change in the area stimulated. Thus, pressures and forces were related by the fixed area stimulated. For convenience, we simply refer to the 3 stimulus magnitudes as force levels. We controlled force levels and timing through custom-built

**Figure 1:**
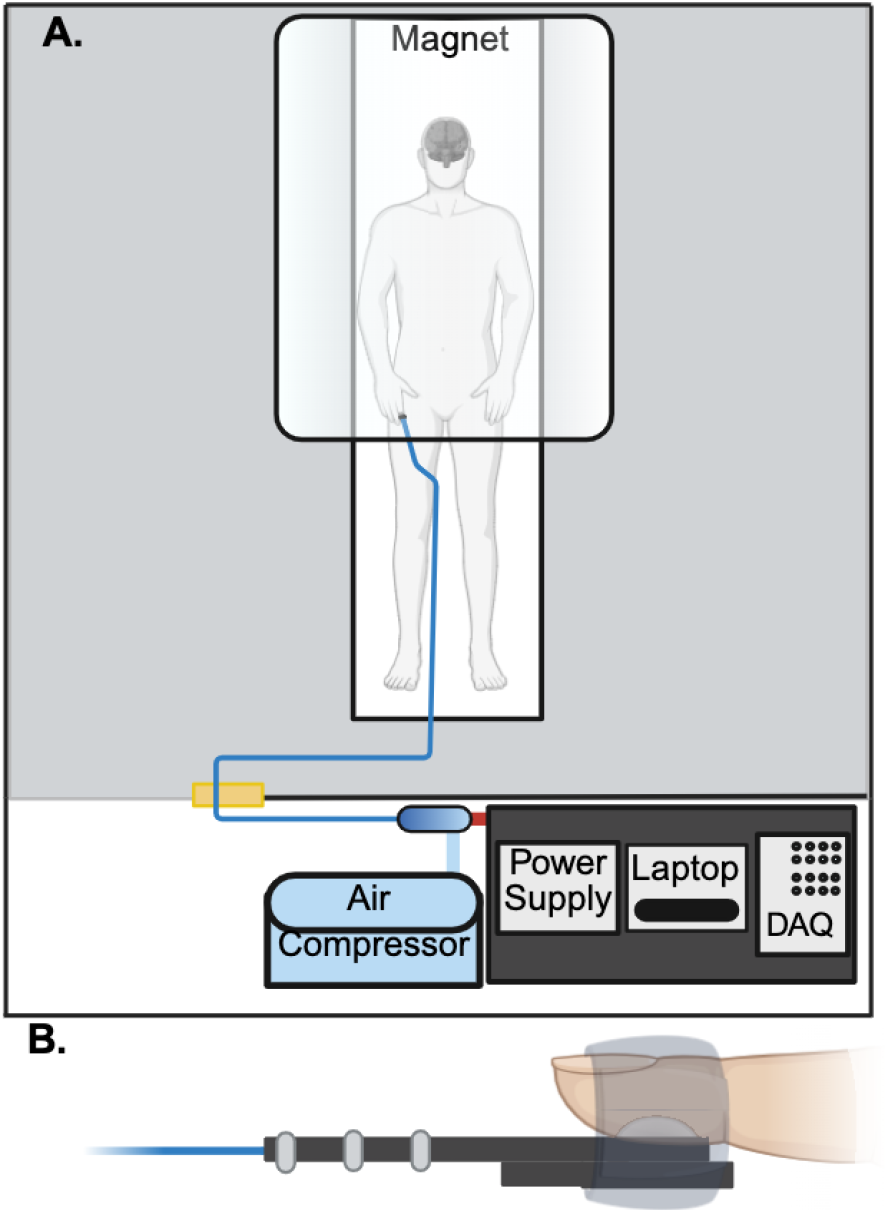
fMRI Compatable Tactile Stimulator: **(A)** Schematic of the tactile stimulator system we used in this fMRI study. All electronic components were housed in the MRI control room. The pneumatic tubing entered the MRI scanner room via a wave guide. All components within the MRI scanner room were non-ferrous (KalantaryArdebily et al., 2024). **(B)** View of the tactile stimulator which was affixed to the participant’s finger using medical tape.

Python-based software. This software is integrated with a data acquisition system (Quanser Q8-USB; Markham, ONTARIO, Canada) to monitor and control pressure (KalantaryArdebily et al., 2024). Stimuli were jittered with interstimulus intervals ranging from 8-12 s (mean:10 s). Stimulus timing was determined using OptimizeX (Spunt, 2016), with each force level as its own condition. Although, stimulus timing and duration were delayed by the pneumatic tubing, this delay was accounted for and verified using the tactile simulator’s pressure recordings.

### 2.3 Functional Magnetic Resonance Imaging

Participants were asked to rest on the MRI table in a head-first supine position with their arms to their sides and their palms lying on their hips (Fig. 1A)^1^. During the scans, participants were instructed to lie still and focus on a visual fixation symbol. They were asked not to cross their arms to avoid tactile perceptual confounds associated with body position (Azañón et al., 2015). We placed the tactile stimulator on the subject’s index finger, centered to match the midline of the distal phalanx. We attached the stimulator using medical tape (Fig. 1B). Data were collected using a 3 T Siemens Prisma with a 32-channel head coil. fMRI scans used a multi-echo EPI pulse sequence provided by the Center for Magnetic Resonance Research (CMRR, Minnesota)(FA=70^°^, repetition time (TR)=2000 ms, slice thickness=4 mm, field of view (FoV)=180 mm, multi-band factor=2, TEe1=13.40,TEe2=39.52,TEe3=65.62 ms) (Moeller et al., 2010; Setsompop et al., 2012). We collected 34 axial slices. The T1 anatomical used the Magnetization-Prepared Rapid Gradient Echo (MPRAGE) pulse sequence (FA=9^°^, TR=2300 ms, slice thickness=1 mm, FoV=256 mm, multi-band factor=1, TE:2.9 ms). All participants receiving a total of 90 stimuli. The tactile stimuli were applied across four scans with a variable number of stimuli in each run. This was done to prevent learning the number of events, since participants were asked to keep a mental count of the number of force stimuli detected during each scan to maintain focus on the task, as done in Hämäläinen et al., 2000. The total number of stimuli differed across the four runs (ranging from 18 to 30) to avoid possible learning confounds. The post-scan questionnaire included Likert scales on body comfort, stimulator comfort, and placement, and weather or not variations in stimulus force were perceived (Fig. 1).

### 2.4 Data Analysis

We used used AFNI and FSL and TEDANA (version 0.0.11; DuPre et al., 2021) to analyze the fMRI data. Preprocessing of the multi-echo data included slice-time correction (3dTshift), rigid-body motion correction to the first time point of the first echo (3dvolreg and 3dAllineate), masking (BET with erosion), and multi-echo combination and de-noising (TEDANA, see next paragraph). The combined echoes were then treated as single-echo data and processed using afni proc.py. The afni proc.py pipeline included EPI alignment to the skull-stripped anatomical, normalization to the Montreal Neurological Institute (MNI) template, spatial smoothing with a 4 mm^2^ FWHM Gaussian kernel, and voxel-wise signal scaling to a mean of zero.

Our TEDANA pipeline closely followed that of Reddy et al., 2024. Specifically, we used TEDANA for T2*-weighted echo combination and independent component analysis (ICA) denoising (DuPre et al., 2021). The data were then dimensionally reduced using echo-time-dependent principal component analysis (PCA) (Y.-O. Li et al., 2007). We manually classified independent components from the ME-ICA based on their likelihood of representing BOLD-related activity. Our manual classification was based on echo-time dependence, spatial distribution, frequency distribution, and temporal noise (Reddy et al., 2024). Subsequently, the non-BOLD ME-ICA components were recombined as a single co-variate to denoise subsequent general linear model (GLM) analyses. Specifically, the sub-matrix of all non-BOLD components was decomposed using PCA and the first eigen-time series was used as a GLM co-variate.

We then performed a GLM analysis that consisted of the stimulus timing, 6 motion regressors, and the non-BOLD eigen-time series co-variate. The three stimulus magnitudes were included as different task conditions, and stimulus onset and duration were convolved with AFNI’s dmBLOCK HRF. We refer to AFNI’s dmBLOCK as the standard HRF, as it was designed to characterize a typical positive BOLD response.

### 2.5 ROI Analysis

Our primary analysis was based on scrutinizing ROIs within S1. Beyond ipsilateral versus contralateral classifications, S1 can be subdivided into areas 1, 2, 3a, and 3b, each of which are cytoarchitecturally and functionally distinct (Kaas, 1983). We compared responses between contralateral and ipsilateral BA1, BA2, and BA3b since these areas are responsible for tactile perception (Rosenthal, 2023), whereas BA3a is related to proprioception and nocioception (Lutz & Bensmaia, 2021; Whitsel et al., 2019). We defined Brodmann areas 1 and 2 using the Brodmann MM3dRm atlas, while BA3b was defined using CA ML 18 MNI (Geyer et al., 2000). To focus on areas of S1 involved in hand/finger representation, analyses of both beta coefficients and time series data were restricted to voxels identified as stimulus-responsive at the group level. To do this we used the intersection between atlas definitions and whole-brain group-level GLM analysis. We applied a group-level whole-brain mask to include only voxels significantly responsive (FDR-corrected p*<*0.05) to the tactile stimuli within each atlas-defined region. These responsive clusters were then segmented according to Brodmann’s area labels.

## 3 Results

### 3.1 Perception of Tactile Stimulus

All participants reported perceiving the tactile stimulus on the first segment of their right index finger (Fig. 2A). The participants’ stimulus counts differed only marginally from the true number of stimuli (0.21±1.05). Although the high, medium, and low force stimuli were intermixed within each run, the stimulus count errors were significantly less than 1/3 (the number of low force level stimuli), suggesting that all 3 stimulus force levels were above the perceptual threshold. No significant difference was found between the subjects’ reported stimulus counts and the actual number of stimuli applied across the runs (Kruskal-Wallis statistic = 4.156, p=0.527), indicating that task attention was consistent throughout the runs (Supplemental Fig.1). After the session, the majority of participants (12/14) reported feeling variations in stimulus force(Fig. 2B).

**Figure 2:**
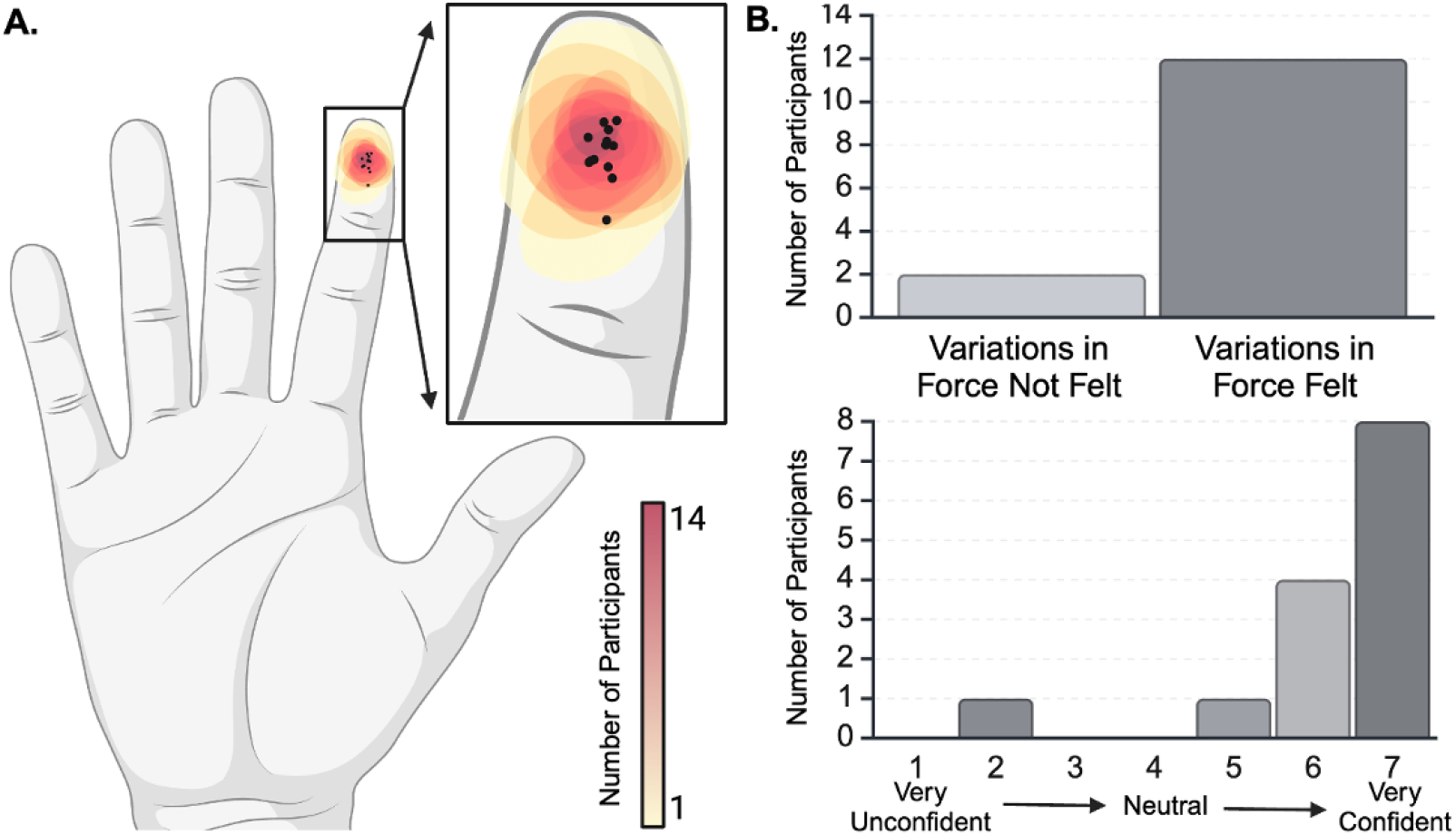
Perception of Tactile Stimulus: **(A)** Heat map of participant-reported stimulus location. Participants were instructed to draw a circle indicating the location and relative size of the tactile stimuli. The black dots represent the centroid of each subject’s reported stimulus presentation area. **(B)** The majority of participants (12/14) noted variations in stimulus magnitude and were confident in their answer.

### 3.2 Distinct Responses in Contralateral vs. Ipsilateral S1

As hypothesized, contralateral S1 had a significant positive response while ipsilateral S1 demonstrated a significant negative response. A repeated measures two-way ANOVA with Tukey correction showed a significant difference between the BOLD response magnitude in the contralateral and ipsilateral hemispheres (non-independent statistics, F(1, 13) = 81.266, p*<*0.0001; Kriegeskorte et al., 2009; Fig. 3). Ipsilateral S1 had a smaller response magnitude across all Brodmann’s areas when compared to the contralateral hemisphere.

**Figure 3:**
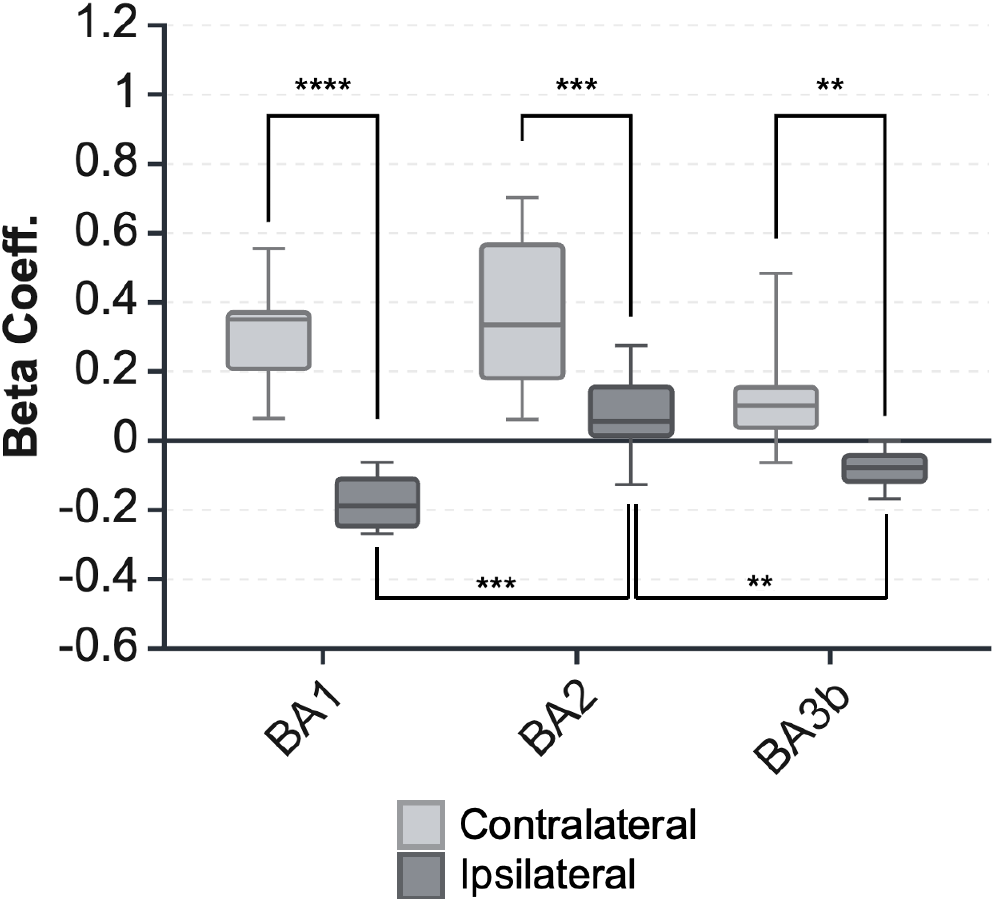
BOLD Response Magnitude in Ipsilateral and Contralateral S1: The BOLD response magnitudes were calculated for each participant within each ROI. Stimulus force-level was used as a parametric modulator, and the resulting beta coeff. represent the mean signal across the 3 force-levels. Error bars represent the 95% confidence interval across all participants. Effect size may be inflated by our ROI definition, which used the intersection of atlas-defined ROIs with whole-brain GLM group results (Kriegeskorte et al., 2009). However, an alternative analysis that avoids this issue, but is less specific to tactile S1, correlates strongly with these results (SFig.2).

### 3.3 Brodmann Area-Specific Responses in Ipsilateral S1

We further hypothesized that within ipsilateral S1, BA1 and BA3b would be the specific subregions with a negative response (Eickhoff et al., 2008; Hlushchuk & Hari, 2006; Klingner, Ebenau, et al., 2011; Klingner et al., 2010, 2014; Schäfer et al., 2012). Testing this, ipsilateral BA1, and BA3b had a significantly lower response magnitude than ipsilateral BA2 (non-independent statistics, p*<*0.0001, p=0.004591 respectively; Kriegeskorte et al., 2009), indicating a non-homogeneous response pattern within the ipsilateral S1 (Fig. 3). Ipsilateral BA2 had a positive response, similar to contralateral S1 regions, with a positive peak around 5 s after stimulus onset (Fig. 4). In contrast, ipsilateral BA1, and to a lesser extent ipsilateral BA3b, had a *negative* peak around 4 s after stimulus onset. We also observed a potential secondary peak around 10 s after stimulus onset (Fig. 4B,F).

**Figure 4:**
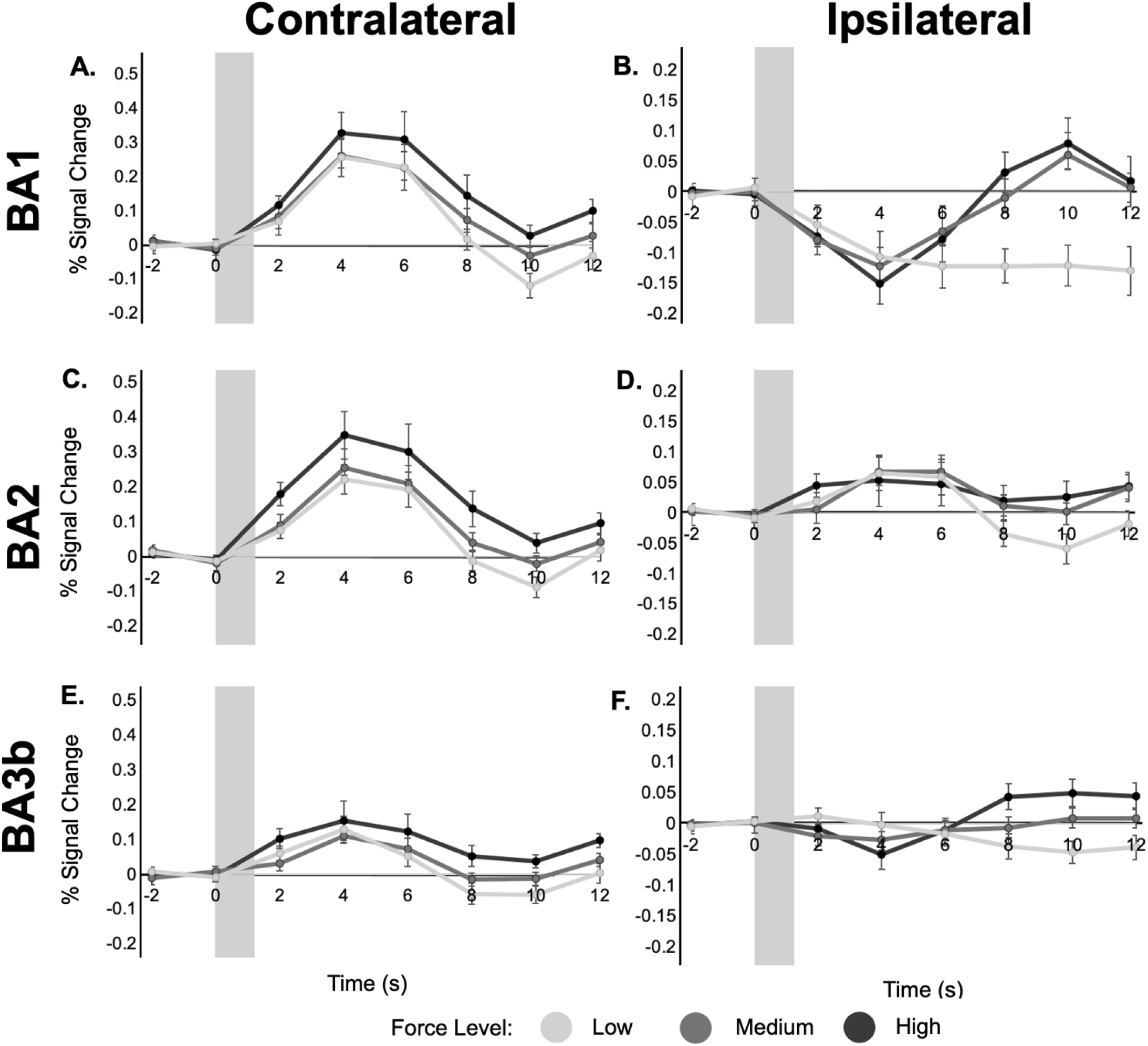
BOLD Response Timeseries in Ipsilateral and Contralateral S1: Each point represents the mean signal change within each ROI, and error bars represent the standard error of the mean. Note the vertical axis scale is fixed in each column, but differs between the contralateral and ipsilateral ROI plots. The gray shaded bar represents the timing and duration of the stimulus.

### 3.4 Ipsilateral BA1 Reflects Biphasic BOLD Activity

Based on Figs. 3 and 4, we further examined ipsilateral BA1. While Fig. 3 showed a robust negative beta coeff., Fig. 4 further suggested that this negative component is followed by a positive overshoot. To assess whether this biphasic response is BOLD-related, we compared percent signal change across echo times, and we calculated the T2* time series as follows:

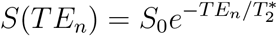

We only examined the high-force level epochs, based on their robust and consistent negative response. Baseline T2* estimates varied across participants, ranging from 46.6 ms to 59.9 ms. Nevertheless, to account for this variability, both percent signal change and T2* data are presented as percent change from baseline (Fig. 5). In ipsilateral BA1, the response magnitude of both the initial negative phase and the secondary positive phase scaled with TE (Fig. 5A). Longer TEs had larger response magnitudes (Fig. 5B). Additionally, we observed a similar biphasic response pattern between the signal change and the T2* timeseries (Fig. 5C). Note that while we focused on the largest force level epochs, we repeated this process for all force levels. For the low force level we observed that the initial negative response exhibited TE-dependence similar to the other force levels. In contrast, the sustained negative signal change plateau observed in response to the low force level stimulus (Fig. 4) was not TE-dependent and a similar plateau was not seen in the T2* response timeseries, indicating that this is likely an artifact.

**Figure 5:**
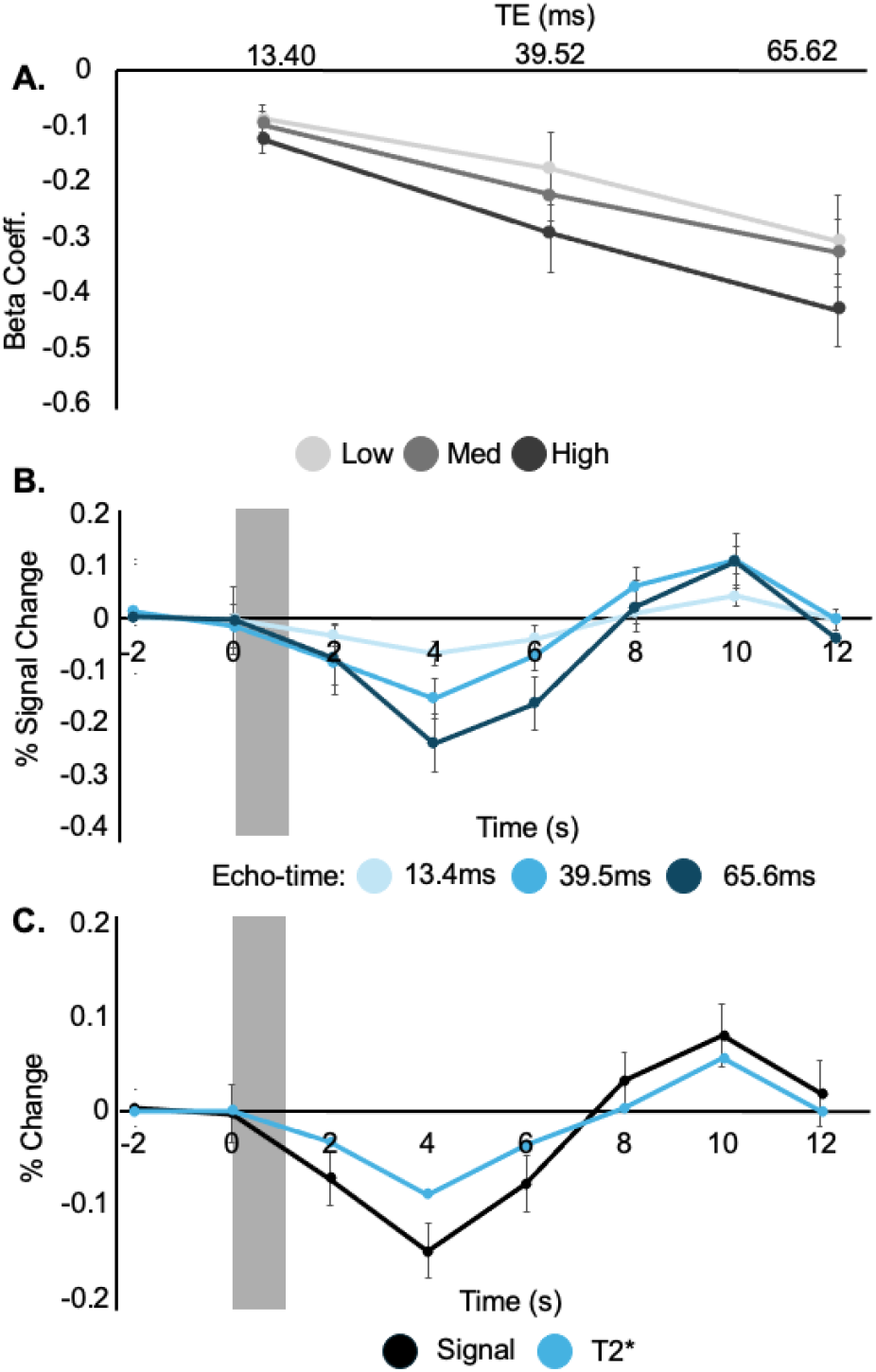
Echo-time dependence in Ipsilateral BA1: **(A)** BOLD response magnitude is larger (more negative) at longer TEs. **(B)** Amplitude is monotonically dependent of echo time with longer echos have greater absolute magnitude change. This indicates that the initial negative response does have echo-time dependence. **(C)** Both the T2* and % signal change time series show an initial negative BOLD response followed by a delayed positive response. The similarity of these two time series suggests that both phases are BOLD-related. Each point represents the mean, and error bars represent the standard error of the mean. All 3 plots include data from only the high force level, which had the largest initial negative and delayed positive responses.

### 3.5 Modeling Biphasic BOLD in Ipsilateral BA1

To model each ROI’s response with a conventional HRF analysis, we estimated the optimal delay for each peak, following methods similar to (Liao et al., 2002). We conducted a delayed HRF analysis to identify the optimal HRF delay that best matched the time series data from these regions. We tested various delay intervals, adjusting the standard HRF onset by 0.5 s steps within a 3 s range around the apparent peak BOLD response, as shown in the time series plots (Fig. 4B). The 6 s window was chosen to capture a broad range of timings without confounding effects from the negative portion of the HRF curve. For each delay, a regression model was applied with stimulus force as a parametric regressor, and the resulting beta coefficients indicated the quality of model fit, with higher values suggesting a better fit. A second-order polynomial curve was then fitted to the group-level data, and the “ideal” delay was defined as its peak (Fig. 6).

**Figure 6:**
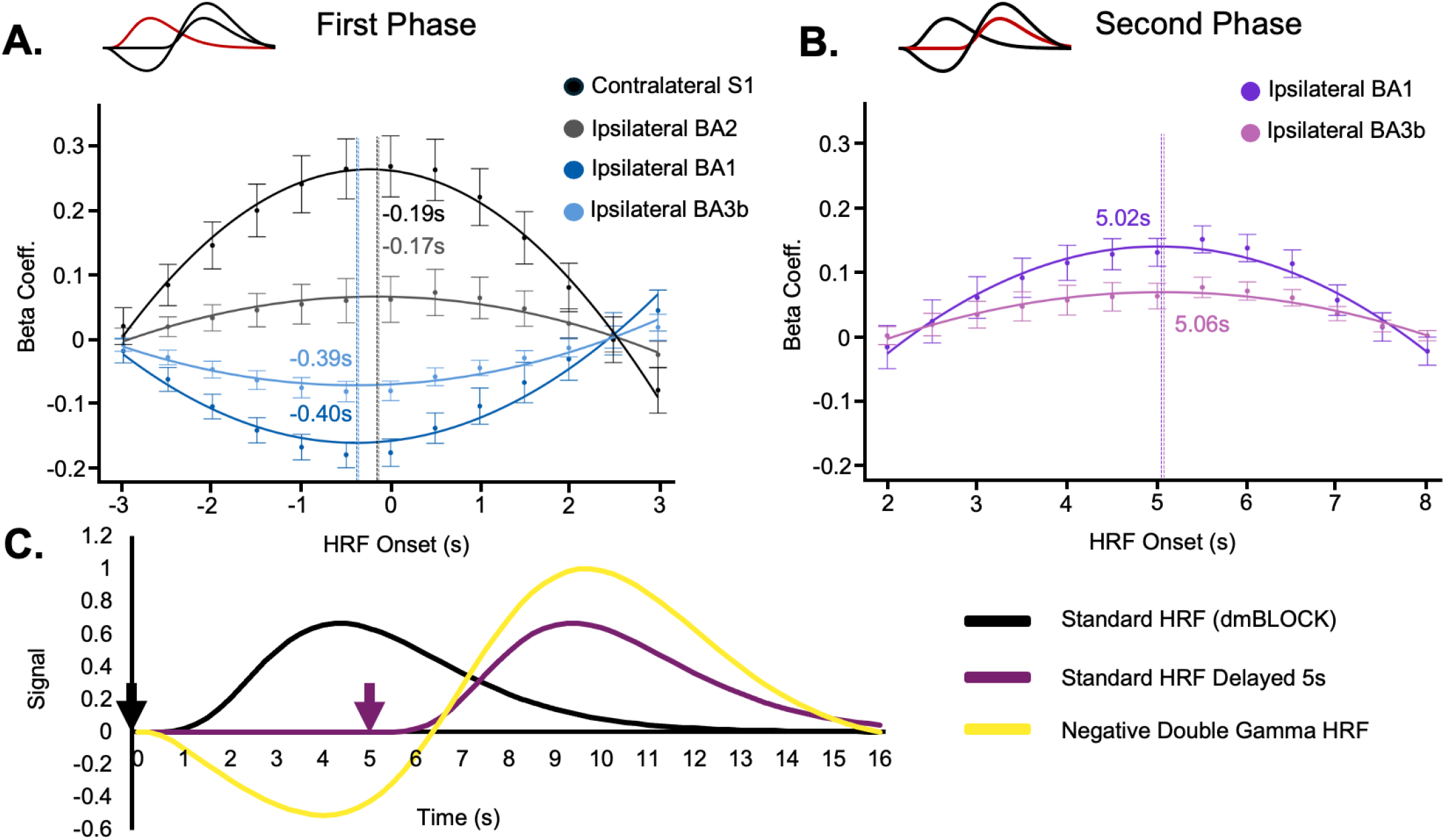
Characterizing the Biphasic Response: **A** and **B** examine the delay in the initial and secondary BOLD response phases, respectively. To do this we preformed GLM analyses (collapsing across all 3 force levels) with a standard HRF (AFNI’s 3dDeconvolve’s dmBLOCK option) across a rang of HRF onset times. **A** and **B** show the GLM fit magnitudes as a function of stimulus timing delay. The resulting beta coeff. vs. delay plots show the mean and standard error of the mean across participants. A higher beta coefficient indicates a better fit between the HRF model and the observed BOLD response. A second-order polynomial was fit to the beta coefficients to estimate the magnitude and delay of the BOLD response (see corresponding polynomial curves). The resulting delay for each ROI is marked with a vertical dotted line. **(A)** Contralateral S1 and ipsilateral BA2 produced positive betas, while ipsilateral BA1 and BA3b produced negative betas. Because we fit these data, we were able to estimate their relative delays (as annotated), however all the delays were approximately zero, especially in the context of a 2 s TR. **(B)** We then characterized the secondary response phase observed in ipsilateral BA1 and BA3b. As shown, the optimal HRF onset was approximately 5 s after the initial stimulus. In other words, if we were to model only the secondary positive response phase in ipsilateral BA1 and BA3b we would stimulus times delayed by approximately 5 s. **(C)** Based on the timing above, a 0 s delay produces the black curve and the 5 s delay produces the orange curve using dmBLOCK. To approximate both these timings, and the negative magnitude of the initial phase in ipsilateral BA1 and BA3b, we tested a negative double gamma HRF, red curve. Arrows indicate the HRF onset time for each model.

In contralateral S1, Klingner, Ebenau, et al., 2011 suggested that negative BOLD may have a different time-to-peak. Thus, they use a standard HFR delayed by 1 s to approximate this negative BOLD response. To quantitatively access any differences in response timing between positive and negative BOLD we include regions with known positive BOLD responses in this analysis. In Fig. 6A, and found that in ipsilateral BA1 the negative peak occurs approximately 0.4 s earlier than the standard HRF would predict. In ipsilateral BA1, the maximal beta coefficient corresponded to a delay of approximately 5 s from stimulus onset (Fig. 6B). Based on this 5 s delay we examined a biphasic HRF using a double gamma approximation with an initial negative phase and a secondary positive phase (Fig. 6C). The 3dDeconvolve negative double GAMMA parameters we used were: TWOGAMpw(9.06,6.06,0.41,5.22,10.31), where the first gamma’s peak and width were 9.06 and 6.06 respectively. The secondary gamma’s peak and width were 5.22 and 10.31 respectively, and the second gamma was scaled by a factor of 0.41.

We then compared a series of GLM maps to assess how each of these HRFs captured the response of ipsilateral S1 (Fig. 7). With the standard (dmBLOCK) model, we see a large negative cluster near ipsilateral (right) BA1. With the negative double gamma HRF approximation, we see a positive cluster in a similar location. This map helps identify the spatial specificity of and corroborates the notion that the secondary positive phase is specific to ipsilateral S1 (Fig. 7B). We see a reduced response area when using the negative double gamma HRF that better resembles the location and shape of BA1, suggesting that the negative double gamma HRF exhibits more specificity than the standard HRF (dmBLOCK). Furthermore, when looking at the distribution of p-values within the ipsilateral S1 cluster, the standard HRF (dmBLOCK) is less concentrated at very low p-values compared to the negative double gamma HRF. This indicates that it is a poorer model of the underlying hemodynamic response within this specific cluster. The improved specificity of the negative double gamma HRF suggests that the true hemodynamic response in this area has a more complex shape than the standard HRF (dmBLOCK) assumes.

**Figure 7:**
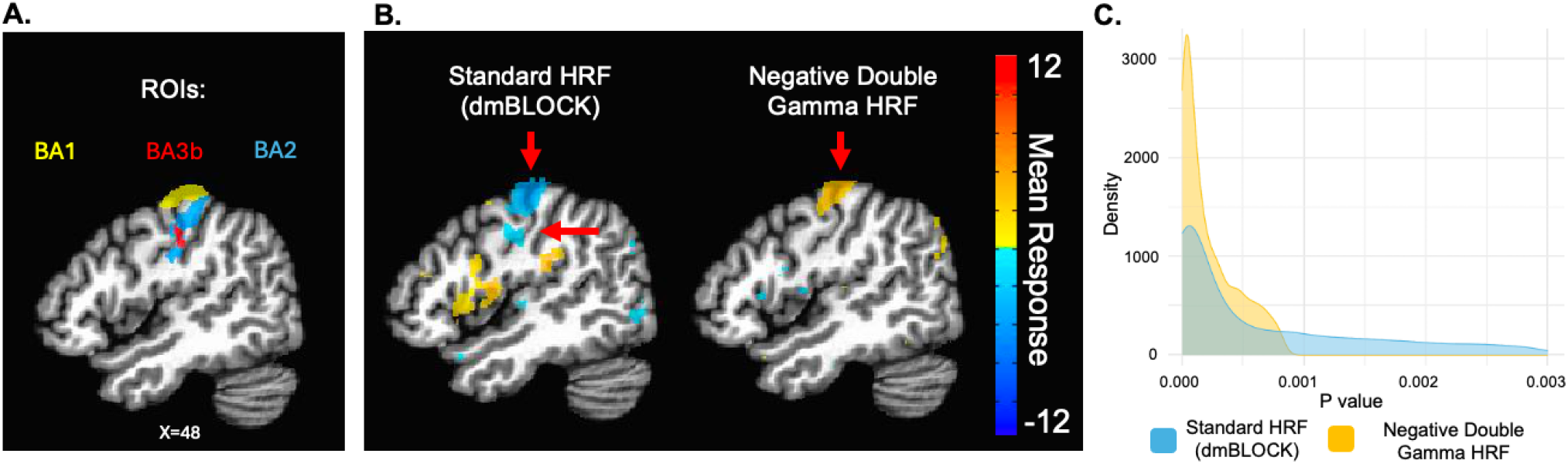
Improved Ipsilateral BA1 Specificity with the Biphasic Model: **(A)** Ipsilateral BA1 and BA2 (defined by the Brodmann MM3dRm atlas) and BA3b (defined by the CA ML 18 MNI). **(B)** Maps generated from the standard HRF (dmBLOCK) and our proposed biphasic negative double gamma HRF from Fig. 6. Arrows indicate GLM results in ipsilateral S1. Note that the sign of the response changes between the standard HRF (dmBLOCK) and the negative double gamma HRF reflecting their negative and positive correlation with the BOLD timeseries respectively. (Maps were thresholded at a voxelwise FDR-corrected p*<*0.05) **(C)** Histograms of p-values within each HRF’s ipsilateral S1 cluster (as highlighted by the red arrows in B), indicating enhanced statistical specificity for ipsilateral BA1.

### 3.6 Force-Dependent Temporal Dynamics

We compared both the standard HRF and negative double gamma HRF to evaluate our ability to detect stimulus force. We performed hemisphere and HRF-specific two-way ANOVAs of region (BA1, BA2, BA3b) and force level (low, medium, high). For the standard HRF we examined both hemispheres. In contralateral S1, we observed a significant main effect of stimulus force level (F(2, 26) = 6.476, p = 0.0052). In contrast, however, the standard HRF did not produce a significant effect of stimulus force in ipsilateral S1 (F(2, 26) = 0.685, p = 0.5127; Fig. 8A). For the negative double gamma HRF, we focused on this ipsilateral hemisphere (where the “negative BOLD response” is relevant). In this case we did observe a significant effect of stimulus force (F(2, 26) = 7.425, p = 0.0028; Fig. 8B). This difference was present even with the inclusion of BA2, which did not exhibit a negative BOLD response in this experiment (Fig. 4D).

**Figure 8:**
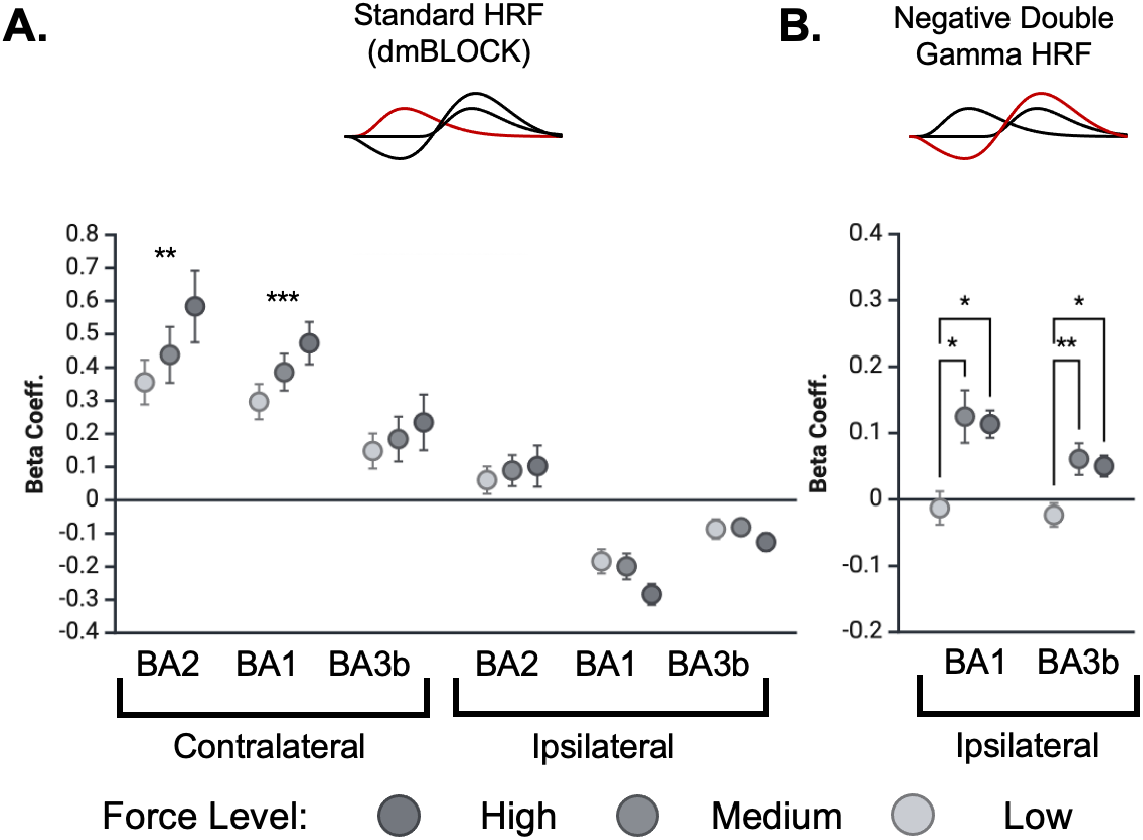
Stimulus Force Modulates Positive and Biphasic BOLD Responses: Shown are the responses for high, medium, and low force levels in S1. Each point represents the mean response magnitude across subjects, and error bars denote the standard error of the mean. **(A)** Shows the standard (dmBLOCK) HRF responses. In contralateral S1 all regions show a positive relationship between stimulus force level and response magnitude. In contrast, none of the ipsilateral regions demonstrated force level dependencies. (one-way ANOVA (F(2, 26); ** p*<*0.01, \*\*\**<*0.001). **(B)** Shows the negative double gamma HRF responses in the ipsilateral S1 regions with negative BOLD. While not as sensitive as contralateral responses, the negative double gamma fits show enhanced sensitivity to force level compared to the ipsilateral fits in A (one-way ANOVA with pairwise comparisons using Tukey’s correction; * p*<*0.05)

As a final test of our hypothesis we compared a biphasic negative BOLD HRF with the stimulus duration found in previously published tactile and visual studies (de la Rosa et al., 2021; Hlushchuk and Hari, 2006; Kastrup et al., 2008; Shmuel et al., 2006; See Fig. 9). As mentioned, most prior reports have used block designs. While the TWOGAMpw parameters are easy to evoke in AFNI and were appropriate for our event-related dataset, this parameterized negative double gamma model produces an exaggerated positive plateau between the initial negative and subsequent positive peaks when convolved with longer stimulus durations. For comparison wit Fig. 9 see supplemental Fig.3. Thus, to compare with those past reports we did not use the general purpose TWOGAMpw approximation, but instead used a fixed-TR, piecewise linear model to produce Fig. 9. Specifically we used a sampling rate of 0.25 s and an HRF with the following parameters in MATLAB ([[0 0] [-.2:-.2:-1.6] [-1.6* ones(1,7)] [-1.5:0.15:1.4] [1.3:-0.2:0.1]]) to conduct a qualitative comparison of those past studies.

**Figure 9:**
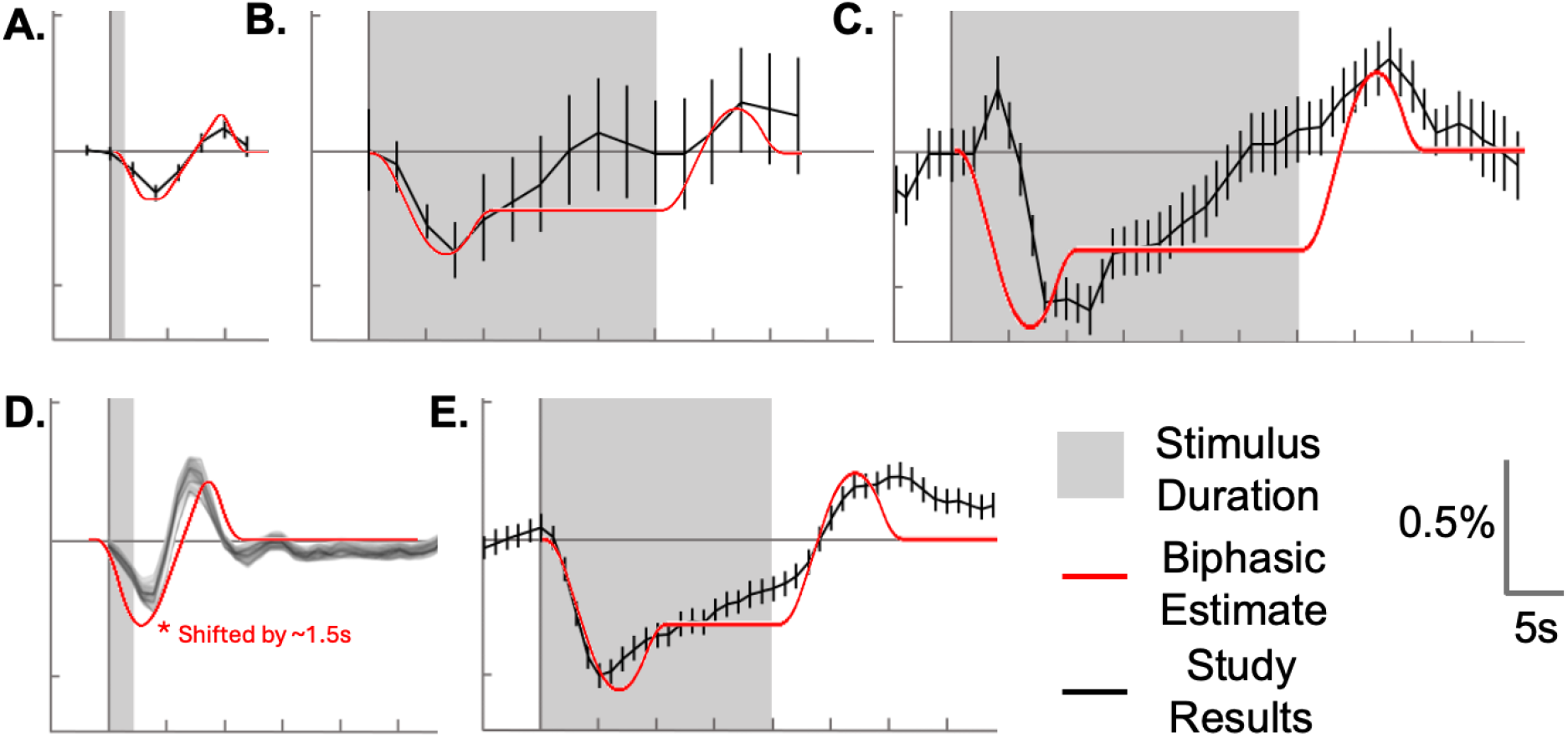
Qualitative Comparison Across Tactile and Visual Studies: A piecewise linear biphasic HRF was convolved with the stimulus duration of prior studies (red curves in A-E). We qualitatively compared this with fMRI signal changes from several studies (black curves in A-E). The studies include tactile stimulation from **(A)** the current study (Fig. 5C), **(B)** Hlushchuk and Hari (2006, Fig.2), and **(C)** Kastrup et al. (2008, Fig.3b). In addition we show responses to visual stimulation from **(D)** de la Rosa et al. (2021, Fig.2), and **(E)** Shmuel et al. (2006, Fig.1d). For the de la Rosa study, our HRF was shifted 1.5 earlier to align with the reported data. (Black curves adapted from the corresponding studies).

Fig. 9A shows our average response to the “high” force-level stimulus in ipsilateral BA1 from Fig. 4B, along with the piecewise linear HRF convolved with our stimulus duration. In Fig. 9B-E we similarly convolved the piecewise linear HRF with the stimulus duration specific to each study. Fig. 9B shows a negative BOLD response elicited in ipsilateral S1 using a 25 s tactile stimulus (Hlushchuk & Hari, 2006). Similarly, Fig. 9C shows an example from ipsilateral S1 from a 30 s block-design electrotactile stimulus to the right median nerve (Kastrup et al., 2008). In visual cortex, Fig. 9D shows results from a short duration (2 s) visual stimulus eliciting responses in ipsilateral primary visual cortex (V1) (de la Rosa et al., 2021) (note in this case we shifted our piecewise linear convolution result by 1.5 s). This 1.5 s dependency may have come from a few sources, including some misunderstanding of stimulus timing on our part. Finally in Fig. 9E we show a 20 s block-design response in peripheral V1 (Shmuel et al., 2006). Some general observations from Fig. 9 include the fact that the initial negative phase of the piecewise linear biphasic HRF aligns well with the time series observed in Hlushchuk and Hari, 2006, de la Rosa et al., 2021, and Shmuel et al., 2006. Despite the 1.5 s dependency the de la Rosa et al., 2021 results nevertheless demonstrate strong similarity between the two waveforms. Notably, our biphasic HRF predicts a delayed positive response in each of these studies that while present, was only noted in the recent de la Rosa et al., 2021 article. Notably, all of these observations hold despite the plateau artifacts in supplemental Fig.3. While qualitative in nature Fig. 9’s cross-study and cross-modal comparisons adds further evidence supporting the plausibility of this study’s biphasic hypothesis.

## 4 Discussion

### 4.1 Discussion of Results

In response to our unilateral tactile stimulus, we observed bilateral responses in primary somatosensory cortex (S1; BA1, BA2, BA3b) and secondary somatosensory cortex (S2). As hypothesized, ipsilateral S1 showed distinct BOLD response patterns. Consistent with prior findings (Eickhoff et al., 2008; Hlushchuk & Hari, 2006; Klingner, Ebenau, et al., 2011; Klingner et al., 2010, 2014; Schäfer et al., 2012), we observed a positive BOLD response in ipsilateral BA2, while ipsilateral BA1 and BA3b exhibited negative BOLD responses.

Extending previous studies, our results and careful review of past reports led us to hypothesize that the ipsilateral negative BOLD response is not simply “negative” as has it been previously characterized. Instead we propose that it is biphasic in nature. As shown in Fig. 4B, the signal in ipsilateral BA1 was not well characterized by a single negative deflection. Instead we observed a biphasic response pattern with a post-negative positive overshoot. To assess whether the biphasic response was truly BOLD-related, we used multi-echo fMRI, examining signal changes across three echo times. This method has been used previously to differentiate BOLD from non-BOLD signal components (Devi et al., 2022; Havlicek et al., 2017; Kundu et al., 2012; N. Li & Jasanoff, 2020). Notably though, this has been done primarily in the context of positive BOLD responses. There remains limited exploration of echo-time dependence in negative BOLD responses, a gap our work begins to address. In ipsilateral BA1, both the negative and positive phases of the signal showed echo-time dependence. We also observed a biphasic T2* response that paralleled the mean signal, reinforcing the interpretation that both components of the biphasic response are BOLD-related and reflect changes in local blood oxygenation (Peltier & Noll, 2002; Serai, 2022).

To model the biphasic response, we used a negative double gamma HRF. Compared to a standard model, the negative double gamma HRF improved stimulus response localization to ipsilateral BA1 (Fig. 7B), and improved GLM statistical specificity in that region (Fig. 7C). Given that the negative double gamma HRF was derived from our data, we further sought to evaluate whether similar patterns generalize to past studies. To do this we convolved the negative double gamma HRF with the stimulus timings used in prior reports. The results in Fig. 9 corroborate a biphasic response was present in both tactile and visual studies. This distinct biphasic response pattern also has been observed and discussed by de la Rosa et al., 2021, where they described an initial negative response followed by a “strong but significantly later positive peak”. Their study used both block design and event-related visual stimuli to examine the negative BOLD HRF. The present study extends this work by framing negative BOLD explicitly as “biphasic,” suggesting parameters to model the HRF, and using multi-echo data to confirm that both phases of the response are BOLD-related.

Our results indicate that the BOLD response to varying stimulus force levels exhibits both spatial and temporal specificity. In contralateral S1, BOLD amplitude was modulated by stimulus force as has been previously reported (Backes et al., 2000). In ipsilateral S1, however, we did not observe force effects with a standard analysis, but amplitude changes were apparent when we modeled the secondary phase of the biphasic response (Fig. 8). Notably, the response magnitude across force levels did not vary monotonically in the ipsilateral cortex (Fig. 4B and Fig. 8B). Instead, we see no response at the low force level and similar response magnitudes at the “medium” and “high” force levels (Fig. 8B). Based on the small range separating “low” vs. “high” force levels, one possible explanation for the similarity between “medium” and “high” is that our current fMRI methods are not sensitive enough to decode differences in these two force levels. On the other hand, we do not think that the similarity in response magnitudes reflects a vascular ceiling effect because “low” did not elicit a response above baseline, and physically and perceptually the difference between “low” and “high” force levels is not large. Nevertheless, these results suggest that there is a minimum stimulus force threshold that needs to be exceed to elicit a biphasic response in ipsilateral BA1 and BA3b. To fully understand the impact of stimulus force in ipsilateral S1, it’s essential to consider the entire biphasic BOLD response, as the initial phase on its own does not differentiate between force levels. It may also be important to examine “negative BOLD” observations in other sensory domains. Analogous to our force-dependent tactile findings, de la Rosa et al., 2021 found that negative BOLD response magnitude varied with visual stimulus features (ie. eccentricity).

Speculating on its functional significance, the early negative phase in BA1 may reflect rapid, stimulus-driven suppression mediated by intrahemispheric inhibitory mechanisms. Given the strong anatomical connectivity between BA2 and BA1, it is possible is that initial excitation in BA2 engages feedforward inhibitory circuits that transiently suppress BA1. This suggests a gating or contrast-enhancement function, momentarily suppressing non-relevant or spatially diffuse signals to enhance the fidelity of tactile representations (Schäfer et al., 2012). Pessimistically, the secondary phase may arise simply from a vascular rebound effect, alternatively it is intriguing to consider the possibility that this positive BOLD signal indicates a delayed recruitment of BA1 into a broader integrative process. If the delayed response we observed is indeed related to neural activity, this would support previous research suggesting that the ipsilateral S1 receives excitatory input from polysynaptic intrahemispheric pathways and/or receives feedback from associative regions such as S2 (Blatow et al., 2007; Chung et al., 2014). This biphasic response profile challenges the traditional view of ipsilateral S1 as passive or purely inhibitory. Instead, it could suggest that ipsilateral BA1 participates in a temporally structured process of tactile encoding, where early inhibition may serve to sharpen response selectivity, (Kastrup et al., 2008; Schäfer et al., 2012), and late excitation may reflect integration of higher-order or bilateral sensory cues (Hämäläinen et al., 2000; Klingner et al., 2016; Tamè et al., 2015).

Continued observations across multiple sensory modalities are needed to more generally relate neural activity to BOLD responses (Nelson & Mayhew, 2025). This not only would aid in improved characterization of the biphasic ipsilateral response but also provide multiple windows to interpret its functional meaning. Previous explicit observations of negative BOLD’s positive overshoot are rare (as an notable exception see de la Rosa et al., 2021). Even more rare has been the attempt to seek mechanistic interpretations, but Gouws et al. (2014) demonstrated that a significant portion of the positive overshoot in visual cortex could be explained by post stimulus eye blinks. Their interpretation was that the positive overshoot was, at least in part, driven by an increase in blinking after stimulus cessation. We hypothesize that currently unobserved analogous behavioral and/or physiological events exist for ipsilateral S1, and for negative BOLD more generally. As an alternative explanation to the Gouws et al. (2014) results, we hypothesize that instead of blinks causing increased overshoot, the intrinsic overshoot component of the biphasic BOLD response could indicate a “release from inhibition” that enables blinking to resume. The increase blink rate could reflect a compensation for the initial suppression of blinks. Although blink rate explained the subsequent positive BOLD phase, further inspection of their results suggest that it also explained the initial negative phase as well.

#### 4.1.1 Limitations and Future Direction

One possible confounding factor complicating previous findings of negative BOLD in ipsilateral S1 is the variety of stimulation methods and stimulus features used. Somatosensory perception involves the complex integration of various stimulus properties, including indentation depth, stimulus frequency, and stimulus modality (eg. brushing, electrotactile, static pressure). Differences in stimulation modality have the possibility of impacting the patterns of ipsilateral S1 activation observed in fMRI studies (Lipton et al., 2006). In this experiment, we used a pressure-based tactile stimulus with a short stimulus duration. This offered the advantage of stimulating mechanoreceptors, in contrast to electrotactile stimuli, which acts on the nerve itself. Despite this limitation, we find that our tactile biphasic HRF generalizes beyond tactile stimuli and is able to approximate negative BOLD responses in the visual cortex.

One limitation of our analysis was our approach to defining S1 ROIs. To obtain an index-finger-specific signal, we used the intersection of atlas-defined ROIs with whole-brain GLM group results. Thus, the effect sizes suggested in Fig. 3 may be inflated (Kriegeskorte et al., 2009). To assess how our approach impacted the results, we also examined the larger, less specific atlas-defined Brodmann’s areas. This control analysis yielded comparable patterns of response across hemispheres and Brodmann’s areas (Supplemental Fig.2). We did not find differences in the beta coefficients (F(1,13) = 2.868, p = 0.1142). However, the overall response magnitudes were attenuated (Supplemental Fig.2). This attenuation likely reflects the exclusion of voxels not specific to index-finger/median nerve stimulation.

While our multi-echo analyses suggested that the biphasic response is BOLD-related, this does not eliminate the possibility of vascular confounds. The BOLD signal is sensitive to a range of hemodynamic variables, including blood flow, volume, and hemoglobin oxygenation (Kim & Ogawa, 2012), which may introduce artifacts independent of neural activity (Harel et al., 2002). Thus, although the T2* and signal time courses agree, additional methods (e.g., concurrent cerebral blood flow measures) could help disentangle vascular and neural signal components.

Regardless of its origins, the biphasic response seems to be a robust, though overlooked, feature of ipsilateral S1 (Fig. 9). While our current study used an event-related design, mapping ipsilateral S1 responses has been a historically challenging goal. Thus, the majority of previous studies have relied on block designs, which are known to elicit more robust BOLD responses. A consequence of this, however, is that the convolution of a block design with an HRF obscures the HRF shape more than an event-related design does. But, as Fig. 9B shows, our proposed biphasic HRF is qualitatively consistent with previous studies in S1 and visual cortex.

## 5 Conclusion

This paper focused on the negative BOLD response in ipsilateral S1, with a primary focus on the more robust BA1. In our ROI analysis, our observations led us to hypothesize that BA1’s response is actually biphasic, particularly for the “medium” and “high” stimulus force levels. By demonstrating echo-time dependence and T2* changes during both phases, we provided further evidence that its response is BOLD-related, and not simply an artifact. Our finding extend prior negative BOLD findings that focused largely on the initial negative response. Finally, the biphasic nature and force-sensitive of ipsilateral S1 suggests it has a more dynamic role in tactile processing than previously recognized.

## 6 Ending Sections

### 6.1 Data and Code Availability

De-identified data and code are available upon reasonable request made to the corresponding author. Data presented in this study can be found on online repositories: Neurovault.

### 6.2 Author Contributions

A.C.F.: Study design, data collection, data analysis, manuscript preparation, software development, writing-review and editing. N.K.: Tactile stimulator design, development, and troubleshooting, and software development, writing-review and editing. N.A.R.: Data analysis, writing-review and editing. R.F.: Data collection, writing-review and editing. J.S.: Data analysis, writing-review and editing. J.L.: Data analysis, data management, writing-review and editing. M.G.B.: Multi-echo fMRI protocol design. N.G.: Study conceptualization, funding, writing-review and editing. S.M.L.: Study design, manuscript preparation, funding, data analysis, writing-review and editing.

### 6.3 Funding

Research was partially supported by the Virginia Tech Institute for Critical Technology and Applied Science Junior Faculty Program (PI: Gurari).

## 6.5 Acknowledgments

We thank the participants for their valuable time and contribution to this study.

## 7 Supplementary Material

**Figure S1:**
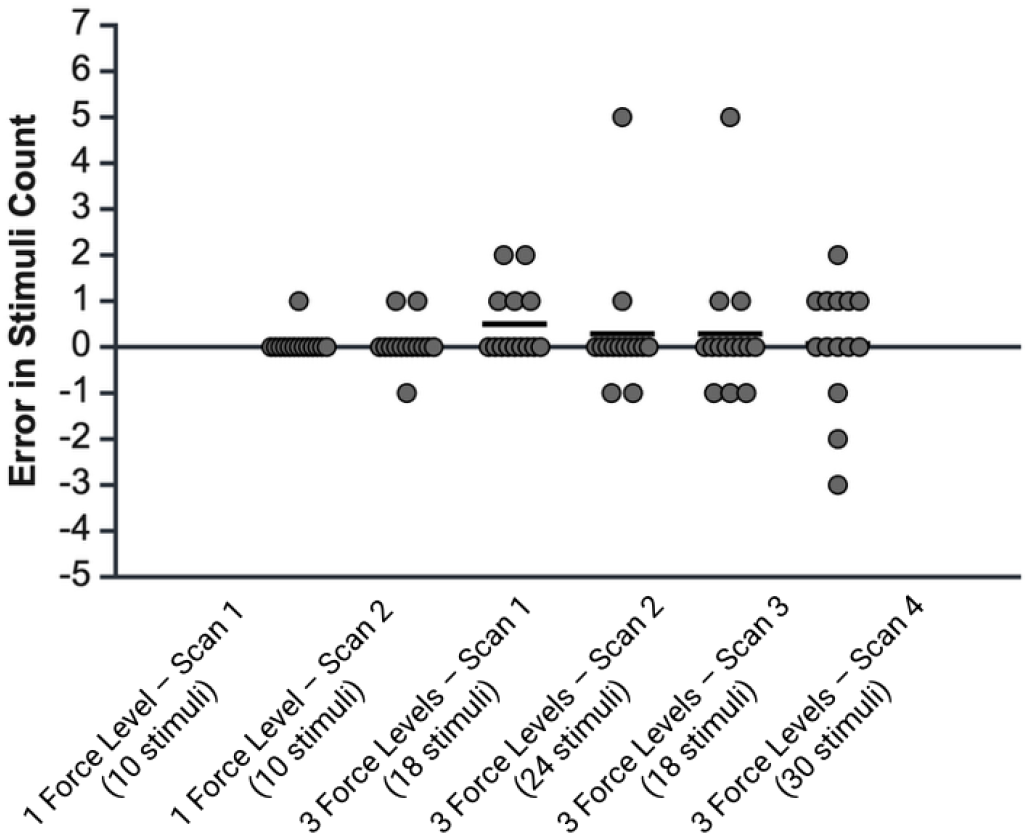
Errors in Subject’s Stimuli Counts: Scans are presented in sequential order from left to right. Each point represents the error in a subject’s reported stimulus count and the true number of stimuli applied. subject counts were never less than two-thirds of the total stimuli applied, indicating that all three stimulus force levels were above the subject’s perceptual thresholds.

**Figure S2:**
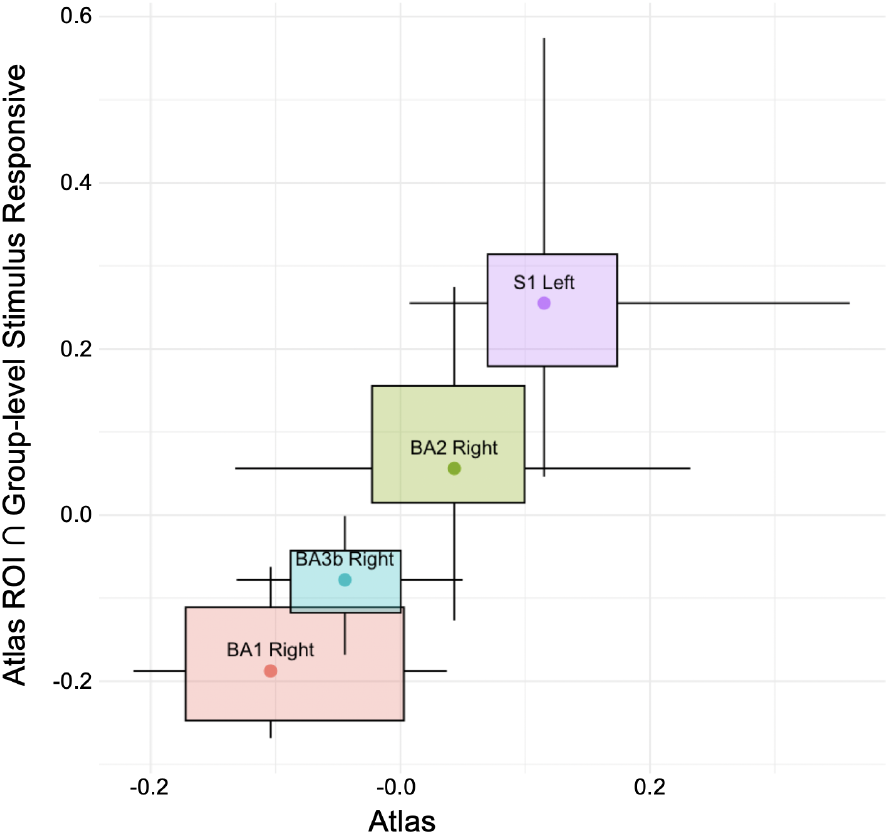
Approaches to Defining ROIs: Box-and-whisker plot examining the beta coeff. between the two approaches for defining our regions of interest within the primary somatosensory cortex. Values represent beta coeff. and the x and y axies correspond to the two approaches to defining ROIs discussed above. This analysis used stimulus force level as a parametric regressor. The error bars represent the 5th and 95th percentile range.

**Figure S3:**
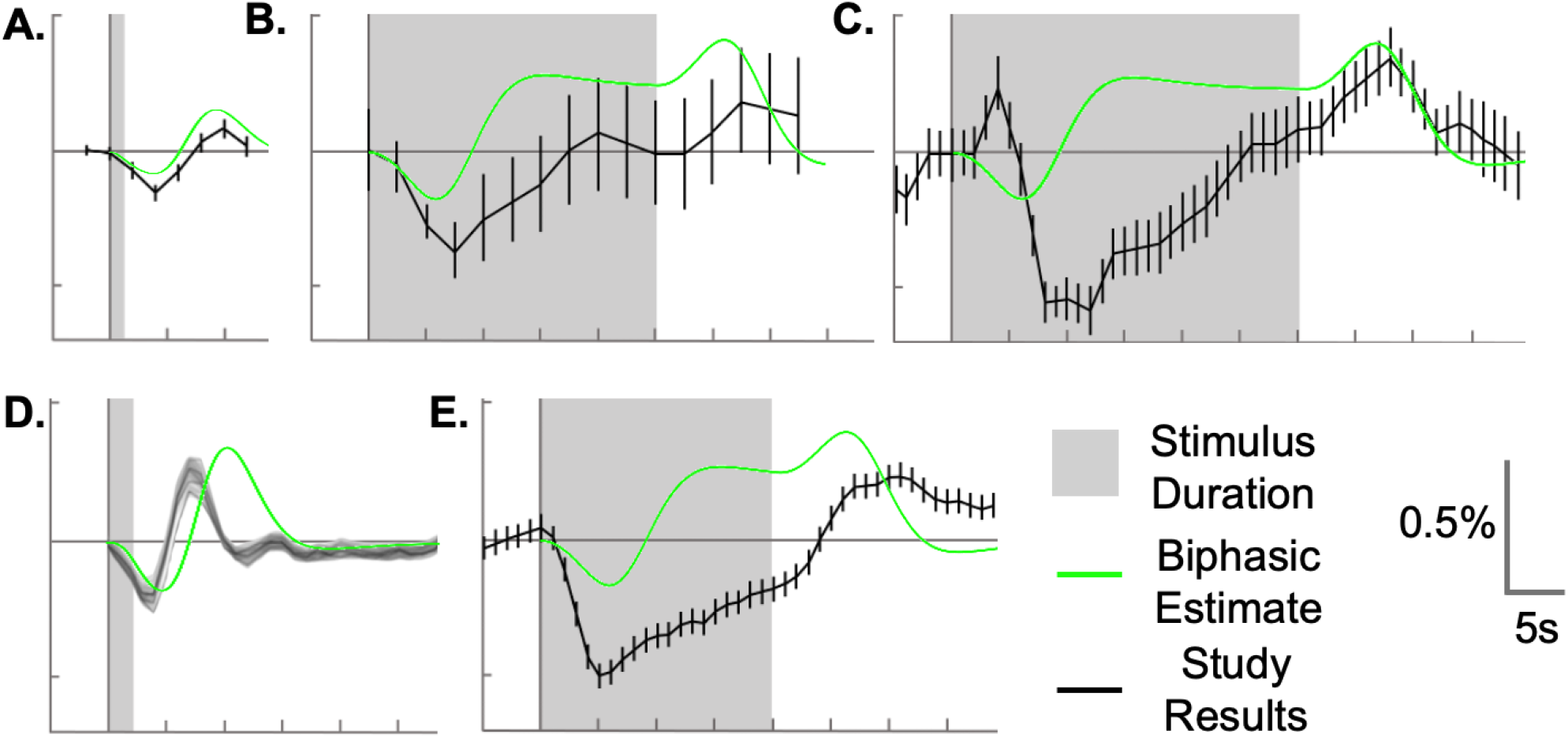
Qualitative Comparison Across Tactile and Visual Studies: The TWOGAM HRF was convolved with the stimulus duration of prior studies (red curves in A-E). We qualitatively compared this with fMRI signal changes from several studies (black curves in A-E). The studies include tactile stimulation from A. the current study (Fig. 5C), **B** Hlushchuk and Hari (2006, Fig.2), and C. Kastrup et al. (2008, Fig.3b). In addition we show responses to visual stimulation from D. de la Rosa et al. (2021, Fig.2), and E. Shmuel et al. (2006, Fig.1d). For the de la Rosa study, our HRF was shifted 1.5 earlier to align with the reported data. (Black curves adapted from the corresponding studies).

## Footnotes

1 Figures were made with BioRender

